## Supplementary material for "*Forget and Forgive*: A Neurocognitive Mechanism for Increased Cooperation During Group Formation": Table S1

**Supplementary Materials**

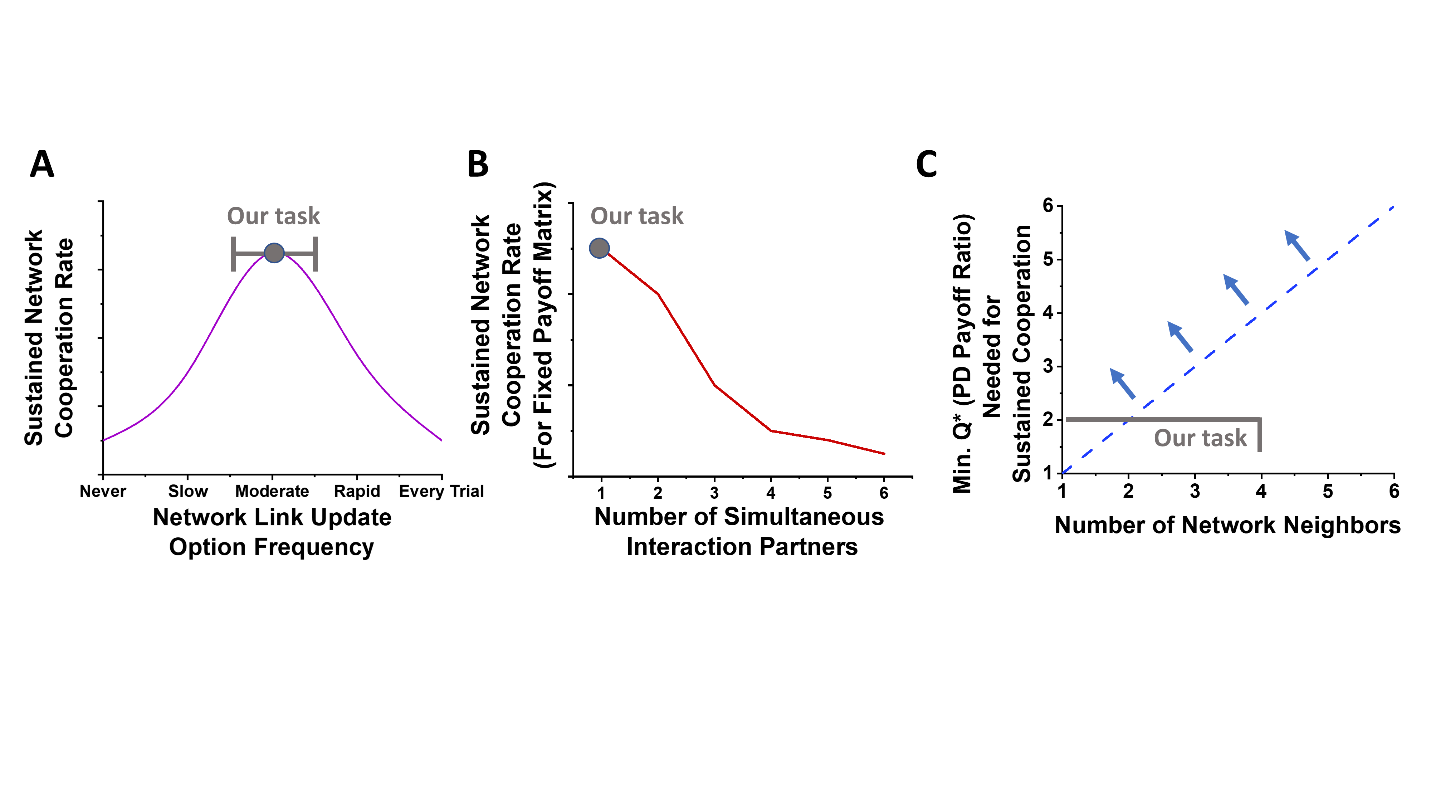

**Figure S1.** **Motivation for iterative embedded-dyad network prisoner’s dilemma design**

(A) Sketch conceptually summarizing prior results suggesting dynamic link variants of network prisoner’s dilemma (PD) foster cooperation in iterative tasks above static link variants, as long as the link update frequency is not too frequent (Rand et al., 2011). We chose a moderate update frequency for both newcomer and link break options (Methods). (B) Sketch conceptually summarizing prior results finding for larger group sizes and fixed payoff matrix, that embedded dyadic PD interactions foster cooperation above simultaneous play interactions with many neighbors at once (Grujić et al., 2012; Hamburger et al., 1975; Yamagishi and Cook, 1993; Yamagishi and Hayashi, 1996). (C) In typical PD notation, our task payoff matrix has T=60, R=30, P=0, and S=-30 (Methods). Rand et al. found the condition for cooperation to succeed in typical static network PD designs is Q* > k, for group size k, and Q*=(P+S-R-T)/(R+S-P-T), which we have plotted (Rand et al., 2014). For our task Q*=(0-30-30-60)/(30-30-0-60) = 2. Our task is not static, so this plot should be interpreted as an approximate guideline. However, we have chosen payoff matrix values on the boundary of this plot to avoid biasing prosocial tendencies with unbalanced payoffs.

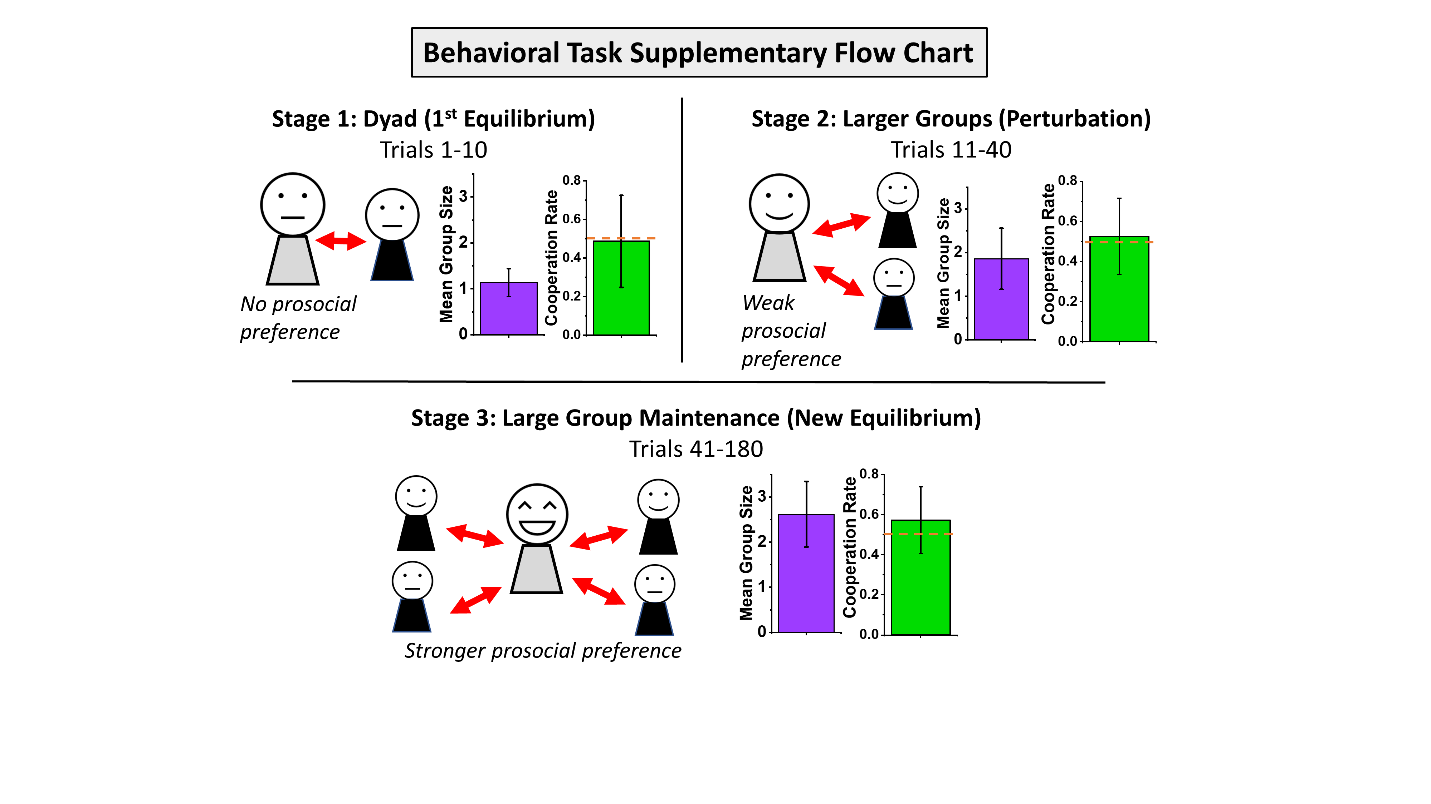

**Figure S2: Additional experimental task flow information and initial cooperation rate results** [related to Figure 1]

Stage 1: For approximately the first ten trials, subjects start in the typical dyad (one partner) prisoner dilemma context. The mean group size is one social partner, and mean cooperation is slightly under 50% for the first ten trials (N=83). Stage 2: Over approximately the next few tens of trials, subjects generally experience larger group sizes of 2-5 social partners, with a mean group size of two social partners and mean cooperation rate of slightly over 50% now for trials 11-40 (N=75). Stage 3: Throughout the rest of the experiment, subjects display more of a “group maintenance mindset” as they dynamically move through the full range of group sizes. The mean group size is near three social partners, and the mean cooperation approaches 60% (N=75). All error bars are SD. The red dotted line on the cooperation rate plots marks 50%.

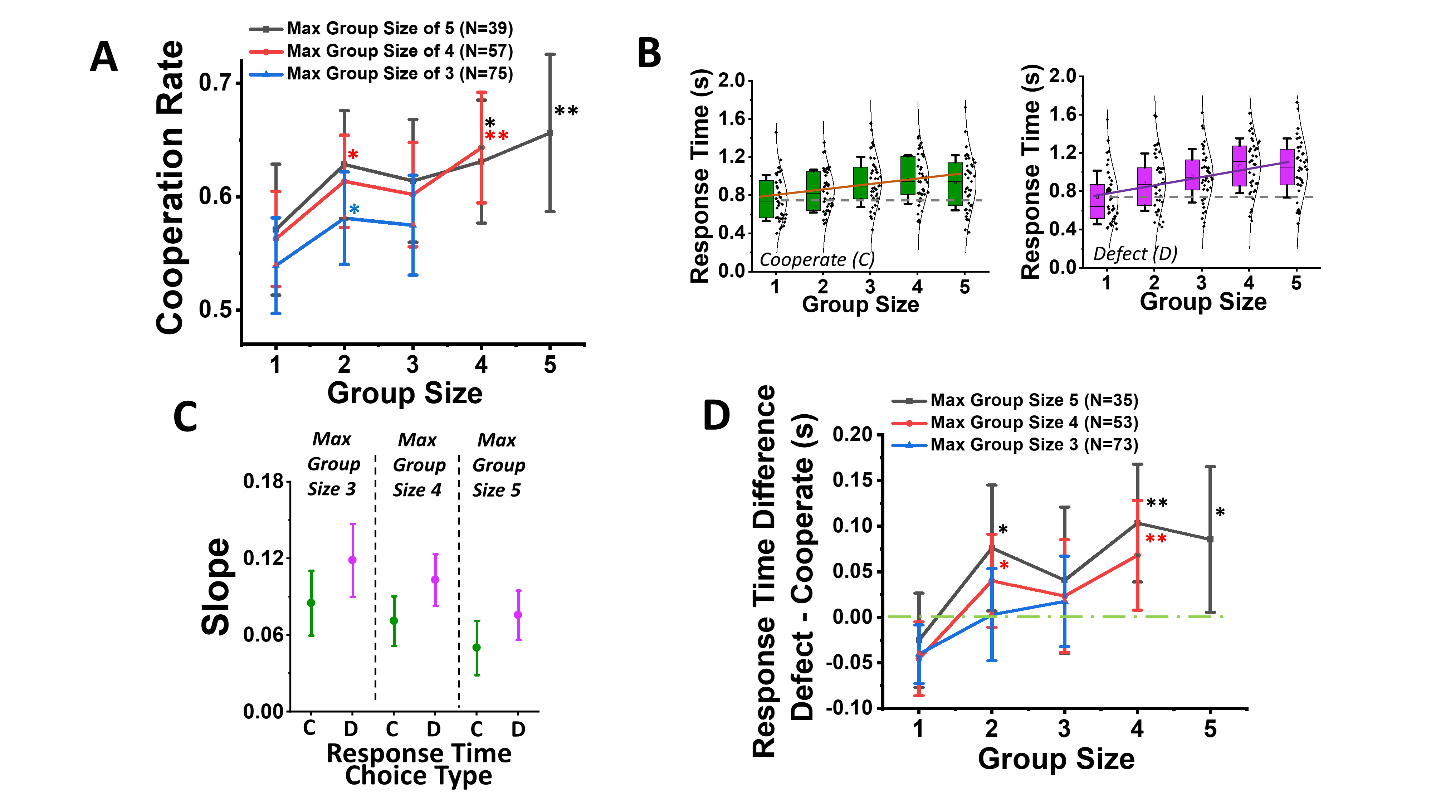

**Figure S3. Group size effects aggregated by maximum group size.**

(A) Between-subject mean cooperation rate (CR) for all subject sampling subsets based on the maximum group size reached in each subject’s session. (B) Plots of the mean response time (RT) per group size, of each subject who had a max group size of 5 (N = 39), as an example to show the underlying distribution of the RT data, with cooperate RTs (left) plotted separately from defect RTs (right). Box plots are 25%/75% box boundaries and 1 SD whisker. Slopes of these plots were calculated for each sampling subset, and plotted in (C) with the same color scheme for cooperation (green) and defect (pink) as (B). Error bars in (C) are 95% CI. (D) RT differences for defect minus cooperate, plotted against group size. In (A) and (D) error bars are 95% CI. Statistical significance was calculated through Holm-Bonferroni-corrected pairwise t-tests relative to the group size of 1, after one-way repeated measures ANOVA, with * being for p < 0.05 and ** for p < 0.01. (Table S1 and S2)

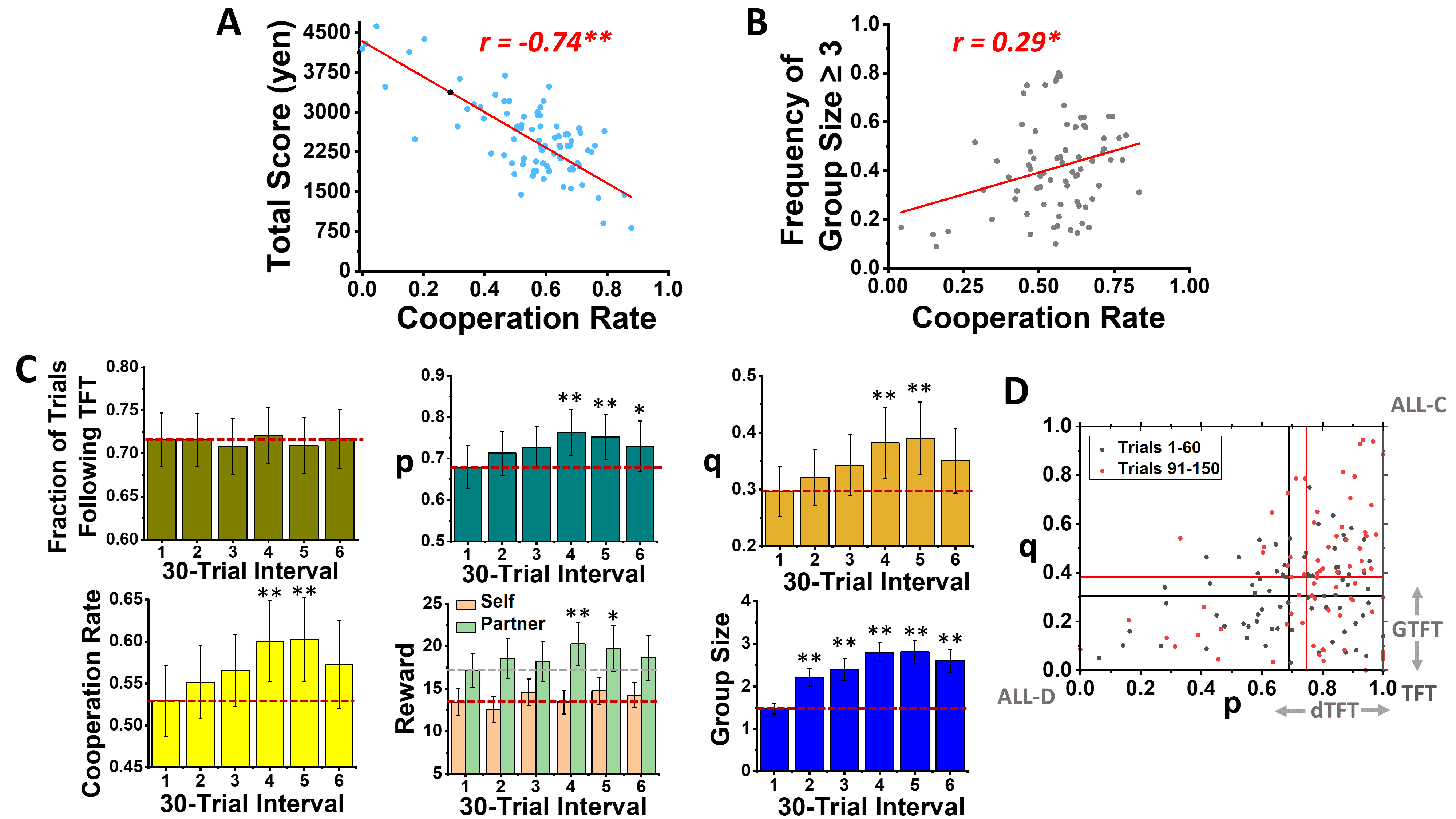

**Figure S4. Subject behavioral strategy analysis**

(A) Cumulative score versus mean cooperation rate for all subjects (N=87, Methods). (B) Plot showing how cooperation rate correlates with the frequency of being in a group size of three or higher (N=75). Pearson correlation coefficients are presented in red text. (C) Plots for 30 trial intervals across the 180-trial session of (left-to-right, top-to-bottom): the mean fraction of trials that subjects’ choices follow pure tit-for-tat (TFT), the mean probability of self-cooperation after a partner cooperation (p), the mean probability of self-cooperation after a partner defection (q), the mean self-cooperation rate, the mean reward rate for the subject (orange) versus partner (green), and the mean group size. Statistical significance was calculated through Holm-Bonferroni-corrected pairwise t-tests relative to first 30 sessions, after ANOVA (N=75) (Table S3). Error bars are 95% CI. (D) Scatterplot of mean q versus p per subject (N=75) for initial trials 1-60 (black) and trials of maximum p-q change 90-150 (red), with mean value per color marked with the straight lines. Typical game theory policies are labeled on the plot for reference: TFT, always cooperate (ALL-C), always defect (ALL-D), generous TFT (GTFT), and our addition, devious TFT (dTFT). * is for p < 0.05 and ** for p < 0.01.

**Table S1: Figure S3a Statistics**

| Pairwise Comparison  (repeated measures, Holm-Bonferroni) | *Prob>\|t\|* (Max Group Size of 3) | *Prob>\|t\|* (Max Group Size of 4) | *Prob>\|t\|* (Max Group Size of 5) |
| --- | --- | --- | --- |
| Group size 1 and 2 | 0.040 | 0.033 | 0.057 |
| Group size 1 and 3 | 0.080 | 0.100 | 0.154 |
| Group size 1 and 4 | **-----** | <0.001 | 0.047 |
| Group size 1 and 5 | **-----** | **-----** | 0.005 |

ANOVA results listed in order of max group size of 3, 4, 5 respectively:

*Mauchly's Test of Sphericity, Prob>ChiSq: <0.001, 0.004, 0.006*

*Mauchly's Test of Sphericity (Greenhouse-Geisser Epsilon): 0.849, 0.820, 0.768*

*Mauchly's Test of Sphericity (Huynh-Feldt Epsilon): 0.867, 0.860, 0.843*

*Repeated measures ANOVA within-subject, Prob>F (Sphericity-Assumed): 0.086, 0.009, 0.072*

*Repeated measures ANOVA within-subject, Prob>F (Greenhouse-Geisser): 0.095, 0.015, 0.091*

*Repeated measures ANOVA within-subject, Prob>F (Huynh-Feldt): 0.094, 0.013, 0.084*

**Table S2: Figure S3d Statistics**

| Pairwise Comparison  (repeated measures, Holm-Bonferroni) | *Prob>\|t\|* (Max Group Size of 3) | *Prob>\|t\|* (Max Group Size of 4) | *Prob>\|t\|* (Max Group Size of 5) |
| --- | --- | --- | --- |
| Group size 1 and 2 | 0.148 | 0.018 | 0.028 |
| Group size 1 and 3 | 0.054 | 0.056 | 0.149 |
| Group size 1 and 4 | **-----** | 0.002 | 0.005 |
| Group size 1 and 5 | **-----** | **-----** | 0.016 |

ANOVA results listed in order of max group size of 3, 4, 5 respectively:

*Mauchly's Test of Sphericity, Prob>ChiSq: 0.989, 0.740, 0.739*

*Mauchly's Test of Sphericity (Greenhouse-Geisser Epsilon): 1.000, 0.968, 0.920*

*Mauchly's Test of Sphericity (Huynh-Feldt Epsilon): 1.000, 1.000, 1.000*

*Repeated measures ANOVA within-subject, Prob>F (Sphericity-Assumed): 0.134, 0.014, 0.045*

*Repeated measures ANOVA within-subject, Prob>F (Greenhouse-Geisser): 0.134, 0.015, 0.050*

*Repeated measures ANOVA within-subject, Prob>F (Huynh-Feldt): 0.134, 0.014, 0.045*

**Table S3: Figure S4c Statistics**

| Pairwise Comparison  (repeated measures, Holm-Bonferroni) | Fraction of Trials Following TFT, *Prob>\|t\|* | “*p*” , *Prob>\|t\|* | “*q*” , *Prob>\|t\|* | Cooperation Rate, *Prob>\|t\|* | Self Reward, *Prob>\|t\|* | Partner Reward, *Prob>\|t\|* | Group Size, *Prob>\|t\|* |
| --- | --- | --- | --- | --- | --- | --- | --- |
| Interval 1 and 2 | 0.993 | 0.181 | 0.401 | 0.349 | 0.390 | 0.223 | <0.001 |
| Interval 1 and 3 | 0.622 | 0.055 | 0.123 | 0.123 | 0.219 | 0.370 | <0.001 |
| Interval 1 and 4 | 0.729 | <0.001 | 0.004 | 0.003 | 0.974 | 0.006 | <0.001 |
| Interval 1 and 5 | 0.649 | 0.004 | 0.002 | 0.002 | 0.158 | 0.025 | <0.001 |
| Interval 1 and 6 | 0.941 | 0.048 | 0.068 | 0.064 | 0.371 | 0.186 | <0.001 |

ANOVA results listed in order of columns (TFT, *p*, *q*, Cooperation Rate, Reward (self), Reward (partner), Group Size):

*Mauchly's Test of Sphericity, Prob>ChiSq:* 0.620, <0.001, <0.001, 0.009, 0.948, 0.006, <0.001

*Mauchly's Test of Sphericity (Greenhouse-Geisser Epsilon):* 0.943, 0.804, 0.815, 0.845, 0.964, 0.836,0.725

*Mauchly's Test of Sphericity (Huynh-Feldt Epsilon):* 1.000, 0.856, 0.868, 0.908, 1.000, 0.892,0.767

*Repeated measures ANOVA within-subject, Prob>F (Sphericity-Assumed):* 0.957, 0.018, 0.015, 0.014, 0.178, 0.089, <0.001

*Repeated measures ANOVA within-subject, Prob>F (Greenhouse-Geisser):* 0.951, 0.027, 0.023, 0.020, 0.181, 0.102, <0.001

*Repeated measures ANOVA within-subject, Prob>F (Huynh-Feldt):* 0.957, 0.024, 0.020, 0.017, 0.178, 0.098, <0.001

**Behavioral patterns for participants with negative social tendency**

Out of 83 participants included in the analysis, 21 exhibited a negative social tendency (*i.e.*, tended to defect more often than cooperate). Analyzing data from only these subjects, revealed no effect of group size on cooperation β = 0.005 95% CI [-0.158, 0.167] p=0.96; but a weak, yet significant negative effect of group size on reciprocity β = -0.075 95% CI [-0.150, 0.001] p=0.046. Similarly, interaction distance did not affect cooperation β = -0.010 95% CI [-0.006, 0.041] p=0.72, but significantly affected reciprocity, β = -0.2501 95% CI [-0.359, -0.141] p<0.001, consistent with the memory effect. Models used were identical for the ones used in the main analysis (Methods). Figure S5 shows these effects.

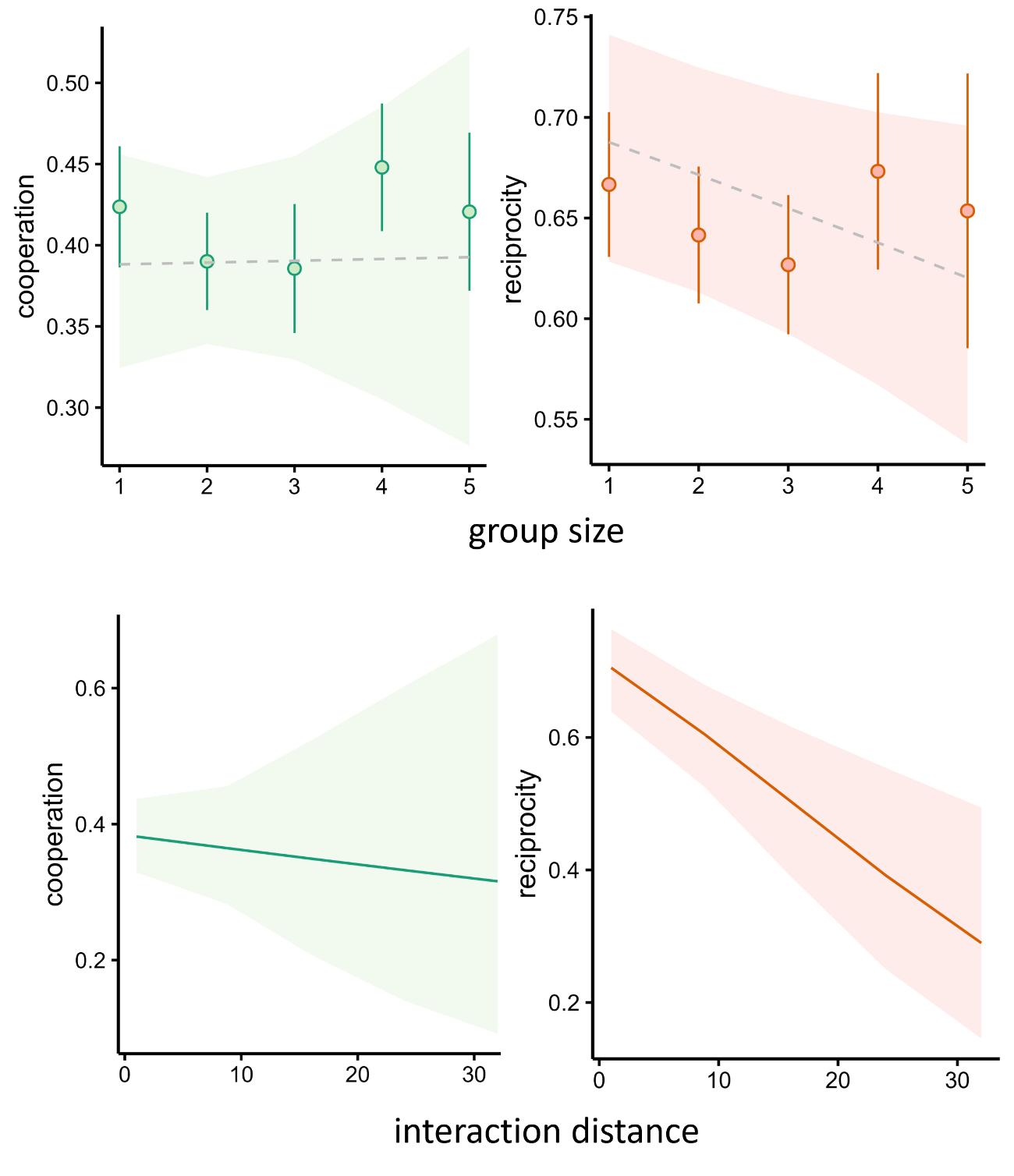
**Figure S5 Behavioral patterns for participants with negative social tendency.** Upper left: cooperation as a function of group size; upper right: reciprocity as a function of group size; lower left: cooperation as a function of interaction distance; lower right: reciprocity as a function of interaction distance. Error bars and ribbons represent 95% CI.

**Recovery of model parameters**

To test recoverability of the parameters of the winning model, we 1) sampled 10 values from posterior distributions of the group-level parameters of the fitted model 2) simulated individual-level parameters from the sampled posteriors 3) simulated experimental data from parameter values sampled in step 2 for 83 synthetic participants, each performing 196 trials of the task 4) Fitted the data simulated in step 3 using the winning model. Figure S6 shows the group-level posterior estimate vs *true* value (simulated in step 2), as well as shows correlations between *true* (step 3) and fitted individual-level parameter values.

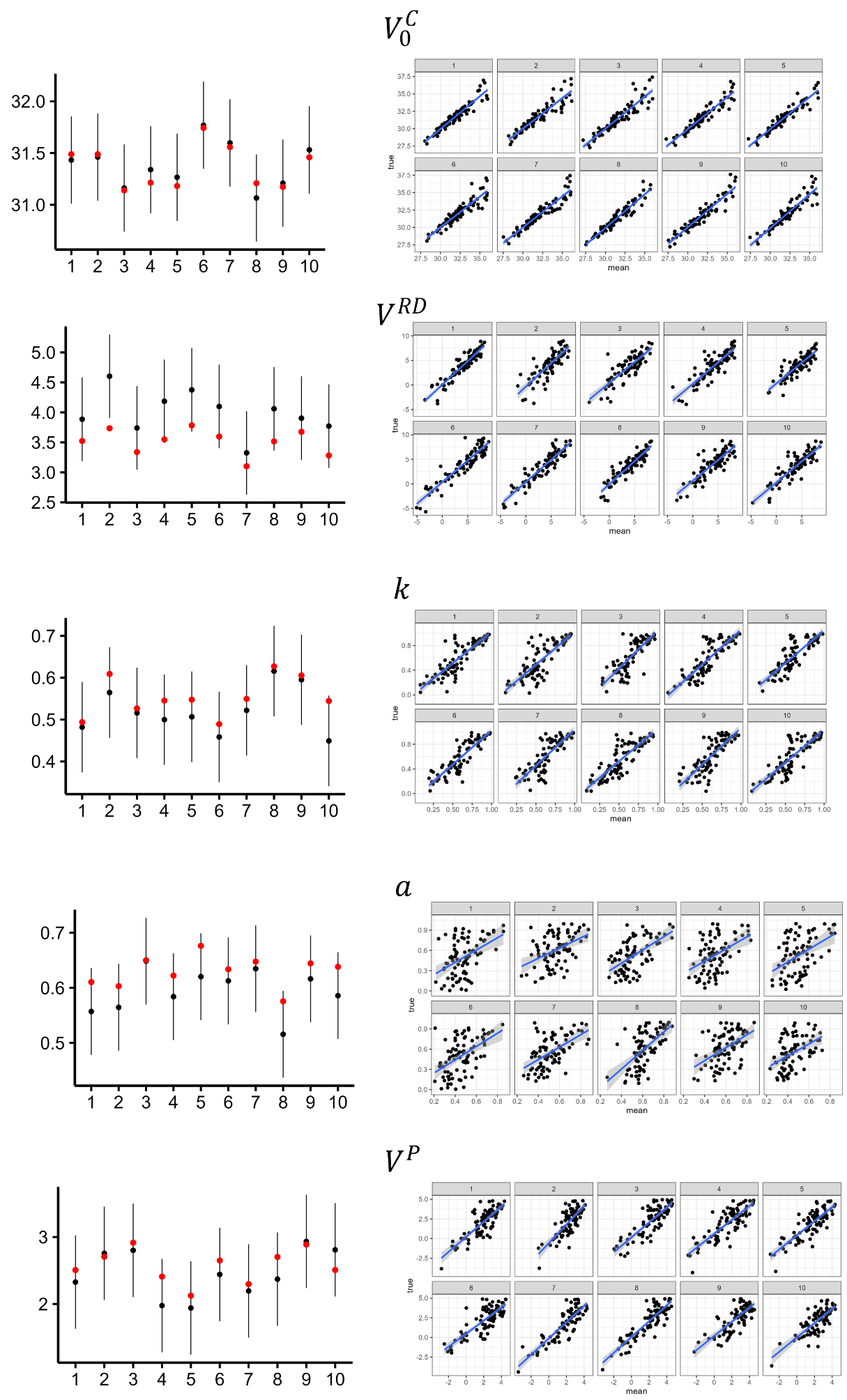

**Figure S6 Parameter recovery.** Each row represents recovery of a given parameter based on 10 simulations. Left column: group-level true values (red dots) vs 90% intervals from posterior model fit. Right column: correlations between true and fitted individual parameters. Each panel represents a single simulation.

**Table S4. Localization of all whole-brain effects**

| Cooperation | | | | |
| --- | --- | --- | --- | --- |
| Structure | Peak MNI coordinates | Cluster size | Peak-level t-value | p(FWE-corr) |
| VS/NAcc | 8 0 -6 | 278 | 6.25 | p=0.003 |
| Reciprocity | | | | |
| Structure | Peak MNI coordinates | Cluster size | Peak-level t-value | p(FWE-corr) |
| dACC | 6 22 44 | 957 | 5.88 | p<0.001 |
| right AI | 24 14 -16 | 527 | 4.74 | p<0.001 |
| left AI | -32 18 -4 | 247 | 4.53 | p=0.005 |
| Forgiveness | | | | |
| Structure | Peak MNI coordinates | Cluster size | Peak-level t-value | p(FWE-corr) |
| right MFG | 32 52 10 | 4829 | 6.82 | p<0.001 |
| left Cerebellum | -49 -56 -32 | 5836 | 6.76 | p<0.001 |
| right AI | 36 18 2 | 636 | 6.03 | p<0.001 |
| left IFG | -46 4 16 | 512 | 5.72 | p<0.001 |
| left SMG | -36 -32 32 | 564 | 5.71 | p<0.001 |
| right SPL | 28 -50 42 | 1814 | 5.63 | p<0.001 |
| left MFG | -22 -2 44 | 206 | 5.63 | p=0.001 |
| left Pallidum | -10 0 0 | 205 | 5.39 | p=0.001 |
| left SPL | -22 -56 34 | 261 | 5.13 | p=0.003 |
| left FFG | -37 -58 -10 | 145 | 4.53 | p=0.047 |
| left MFG | -40 44 6 | 188 | 3.78 | p=0.016 |
| Betrayal | | | | |
| Structure | Peak MNI coordinates | Cluster size | Peak-level t-value | p(FWE-corr) |
| right DLPFC | 40 48 24 | 333 | 6.42 | p=0.002 |
| left DLPFC | -36 40 14 | 185 | 6.35 | p=0.032 |
| right Cerebellum | 34 -60 -48 | 168 | 4.23 | p=0.046 |
| Forgetting | | | | |
| Structure | MNI coordinates | Cluster size | Peak-level t-value | p(FWE-corr) |
| left FFG | 40 -52 -22 | 268 | 5.80 | p<0.001 |
| precuneus | -12 -66 30 | 2177 | 5.52 | p<0.001 |
| right FFG | -38 -48 -22 | 299 | 5.42 | p=0.001 |
| right Cerebellum | 10 -76 -38 | 142 | 5.36 | p=0.047 |

**Supplementary References**

Grujić, J., Eke, B., Cabrales, A., Cuesta, J.A., Sánchez, A., 2012. Three is a crowd in iterated prisoner’s dilemmas: experimental evidence on reciprocal behavior. Scientific Reports 2, 638. https://doi.org/10.1038/srep00638

Hamburger, H., Guyer, M., Fox, J., 1975. Group Size and Cooperation. Journal of Conflict Resolution 19, 503–531. https://doi.org/10.1177/002200277501900307

Rand, D.G., Arbesman, S., Christakis, N.A., 2011. Dynamic social networks promote cooperation in experiments with humans. PNAS 108, 19193–19198. https://doi.org/10.1073/pnas.1108243108

Rand, D.G., Nowak, M.A., Fowler, J.H., Christakis, N.A., 2014. Static network structure can stabilize human cooperation. PNAS 111, 17093–17098. https://doi.org/10.1073/pnas.1400406111

Yamagishi, T., Cook, K.S., 1993. Generalized Exchange and Social Dilemmas. Social Psychology Quarterly 56, 235–248. https://doi.org/10.2307/2786661

Yamagishi, T., Hayashi, N., 1996. Selective Play: Social Embeddedness of Social Dilemmas, in: Liebrand, W.B.G., Messick, D.M. (Eds.), Frontiers in Social Dilemmas Research. Springer Berlin Heidelberg, pp. 363–384.
